# Large-scale single-neuron recording using the uFINE array in human cortex

**DOI:** 10.1101/2025.04.14.648657

**Authors:** Shun Wu, Zhiqiang Yan, Cen Kong, Xia Li, Qiufeng Dong, Youkun Qian, Guangyuan Chen, Beibei Chen, Chi Ren, Junfeng Lu, Xiaofan Jiang, Zhengtuo Zhao, Xue Li

**Affiliations:** Institute of Neuroscience, Center for Excellence in Brain Science and Intelligence Technology, Chinese Academy of Sciences, Shanghai, 200031, China; Department of Neurosurgery, Xijing Hospital, Fourth Military Medical University, Xi’an, 710032, China; University of Chinese Academy of Sciences, Beijing, 100049, China; Department of Neurosurgery, Huashan Hospital, Shanghai Medical College, Fudan University, Shanghai, 200040, China

**Author notes:** These authors contributed equally (Shun Wu, Zhiqiang Yan, Cen Kong).

**Keywords:** Ultraflexible neural electrode, human intraoperative recording, stable single-neuron recording, brain-machine interface

## Abstract

Monitoring neural population activity at single-cell resolution is essential for driving fundamental research and clinical innovations. However, translating advanced recording techniques from animal models to humans remains a significant challenge. Flexible neural electrodes have recently emerged as powerful tools for large-scale single-unit recordings due to their superior biocompatibility and high recording density. Here, we demonstrate reliable, high-density single-unit recordings during intraoperative procedures in human patients using **u**ltra-**F**lexible **I**mplantable **N**eural **E**lectrode (uFINE) arrays. The uFINE array exhibited sufficient mechanical robustness to maintain structural integrity throughout surgical operations. We successfully recorded 616 single units from 10 patients, with up to 135 single units simultaneously recorded. The flexibility of uFINE array minimized signal disturbances from brain pulsations, enabling stable and continuous single-unit detection. Stimulus and response tuning were observed at the level of individual neurons in awake patients. This uFINE-based recording approach offers unique opportunities to investigate human-specific cognitive functions and develop next-generation brain-machine interfaces.

## INTRODUCTION

Single-unit recordings have long been a cornerstone of neuroscience research, offering direct observation of individual neuron activity with exceptional temporal resolution. In the human brain, single-unit recordings provide crucial insights into the neural mechanisms underlying sensory, motor, and particularly human-specific cognitive processes, with broad applications in both basic neuroscience and clinical practice^1,2^.

Traditional approaches to intraoperative single-unit recordings in the human brain, however, are often limited by the number of neurons that can be recorded simultaneously. Commonly used electrode types, such as metal microelectrodes^3,4^ and microwire bundles^5–7^, typically yield a small number of single units per electrode (8 microwires)^8–11^. Although these methods have been instrumental in characterizing the properties of individual neurons, they fall short of capturing the dynamics of neuronal populations. The 96-channel Utah array, which allows for simultaneous recordings of a larger number of neurons, has been widely used in brain-machine interfaces^12–17^. However, its implantation often leads to significant acute tissue damage, making it unsuitable for intraoperative settings that require minimal tissue disruption (for an exception, see Ref^18^.).

Recently, the intraoperative use of Neuropixels probes has significantly expanded the scale of single-unit recordings in the human brain, providing deeper insights into the neural mechanisms underlying complex functions such as language processing^19–23^. However, this silicon-based probe still faces challenges from substantial brain movements during surgery, which can compromise both recording stability and single-unit yield^19,20,24^. Additionally, large lateral brain movements increase the risk of probe breakage, raising concerns about safety and reliability of Neuropixels probes for intraoperative use, as well as their potential for long-term implantation. Thus, there remains a critical need for safer, more effective tools for large-scale single-unit recordings in the human brain.

Ultra-flexible microelectrodes have emerged as a promising solution to these challenges, owing to their excellent biocompatibility and minimal tissue damage^25–27^. These electrodes have demonstrated success in achieving large-scale, long-term stable single-unit recordings in rodents and non-human primates^28–35^. In this study, we present a methodology that utilizes an **u**ltra-**F**lexible **I**mplantable **N**eural **E**lectrode (uFINE) array to record single-unit activity cross cortical depths in the human brain during intraoperative procedures. The flexibility of the uFINE array allows for stable, high-quality recordings even in the presence of rapid brain movements, resulting in high single-unit yields. Using this approach, we also reveal the heterogeneous responses of individual neurons to stimulus and the response period in the dorsolateral prefrontal cortex (dlPFC). Taken together, these results demonstrate the potential of the uFINE array as a powerful tool for intraoperative neurophysiological investigations of the human brain, with applications in both healthy and diseased states.

## RESULTS

### Implantation of the uFINE array in human cortex

Intraoperative single-unit recordings using the uFINE array were attempted in 15 awake patients undergoing Deep Brain Stimulation (DBS) lead implantation or brain tumor resection (Table S1). Successful recordings of single-unit activity were achieved in 10 patients (Figure 1A), while failures in the remaining 5 patients, during the initial troubleshooting stage at each clinical center, were mainly due to excessive electrical noise in the operating room environment and damaged tips of shuttle wires that led to insertion failure. We found that the root mean square (RMS) value of electrical noise could be reduced to 5-10 µV by (1) separating ground and reference (e.g., one placed in the scalp and the other in the cerebrospinal fluid); and (2) turning off the surgical lights and navigator during recordings (Methods). The assembled uFINE array (Figure S2A and methods) and the headstage were connected and encapsulated within a waterproof protective case, forming an enclosed uFINE module (Figure 1B). Furthermore, to prevent damage to shuttle wires tips, we designed a removable protective cap for array tips (Figure S2A) and confirmed the insertion site using a dummy uFINE module before formal implantation (Methods). The 5 patients without single-unit detection were excluded from further analyses in this paper.

**Figure 1.**
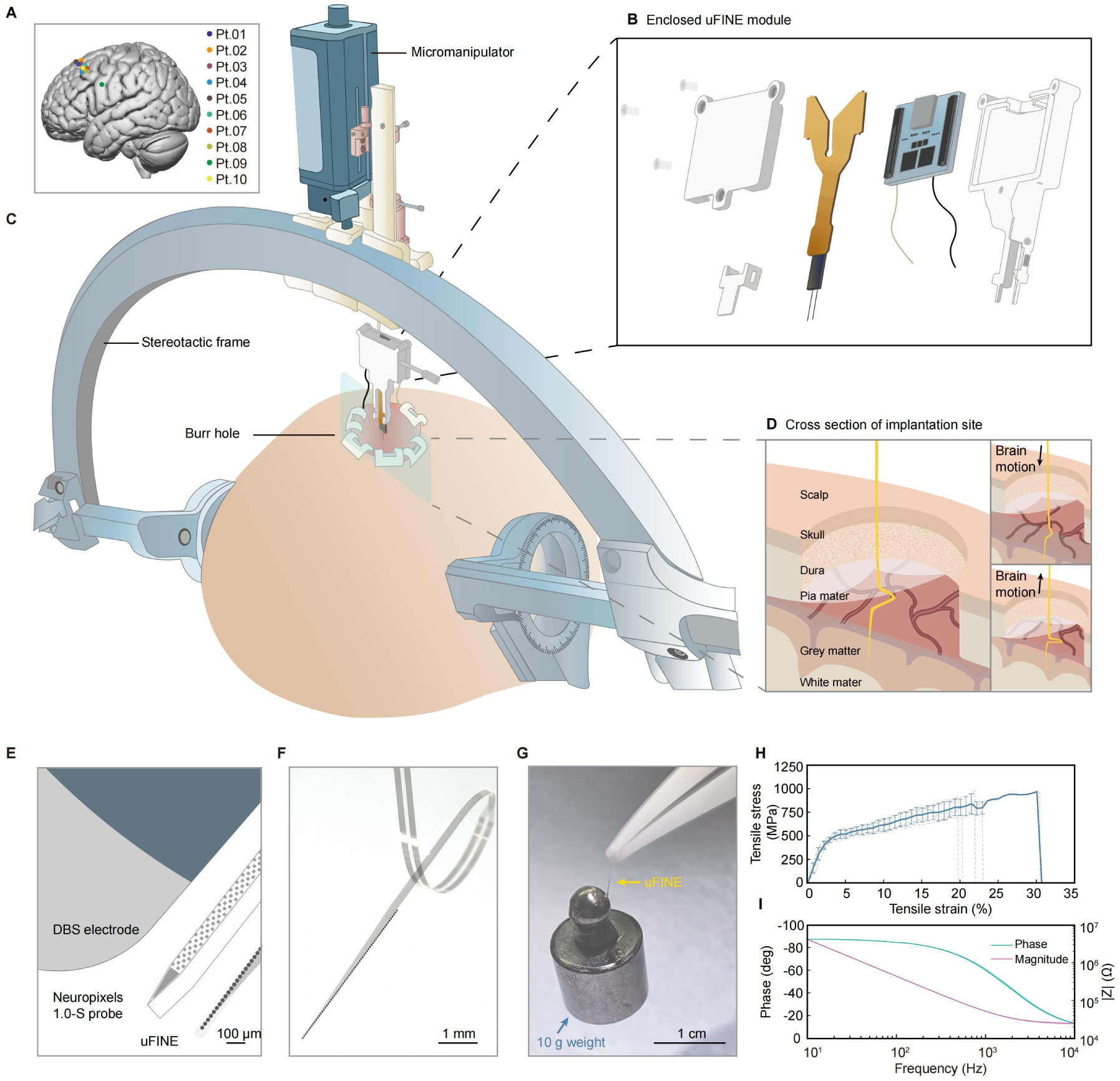
Intraoperative recording using the uFINE array in human cortex. (A) Insertion sites of 10 patients with single-unit detection. (B) Schematic diagram of the enclosed uFINE module. (C) Schematic diagram showing the implantation setup for the uFINE array during DBS surgery. (D) Cross-sectional view of the implantation site showing the flexible uFINE array moving with brain pulsations. (E) Comparison of dimensions between the DBS lead, the Neuropixels 1.0-S probe, and the uFINE array at the same scale. (F) Photo of one uFINE array shank immersed in water, showing its flexibility. (G) Photo of one uFINE array shank holding a 10-gram weight, showing its mechanical strength. (H) Averaged stress-strain curve of the uFINE array shank (n = 5), with gray dashed lines representing the results of individual samples. Error bars, std. (I) Averaged Electrochemical Impedance Spectroscopy (EIS) curve of the uFINE recording sites (n = 6). Shaded areas, std.

**Figure S1.**
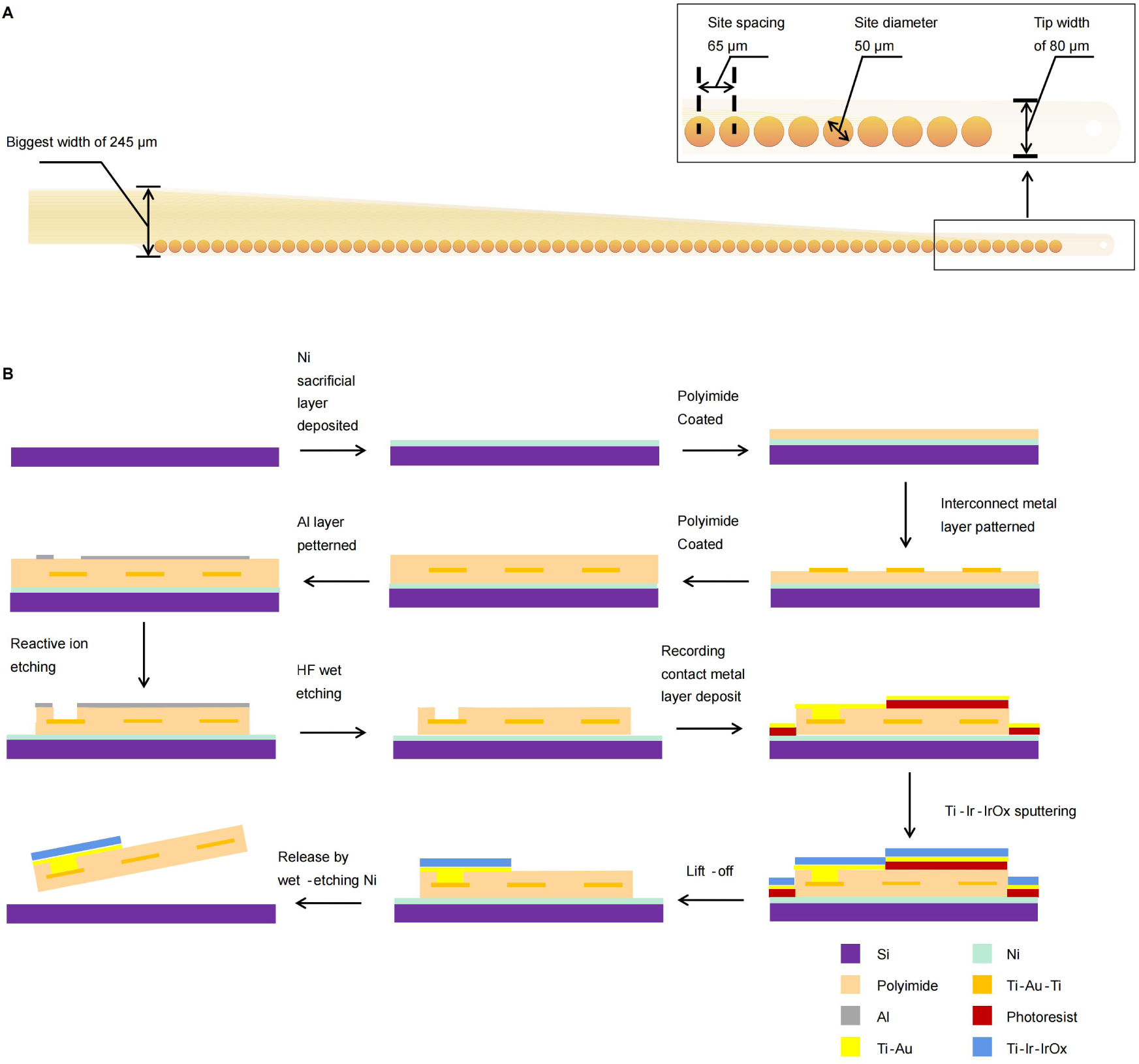
Design and overall fabrication schematic processing of uFINE array for intraoperative use. (A) Schematic diagram showing the shank tip with recording sites. Zoomed-in view shows the dimensions of the recording site, the site spacing, and the shank tip width. (B) Schematics diagrams showing the fabrication steps of the uFINE array (see Methods for details).

**Figure S2.**
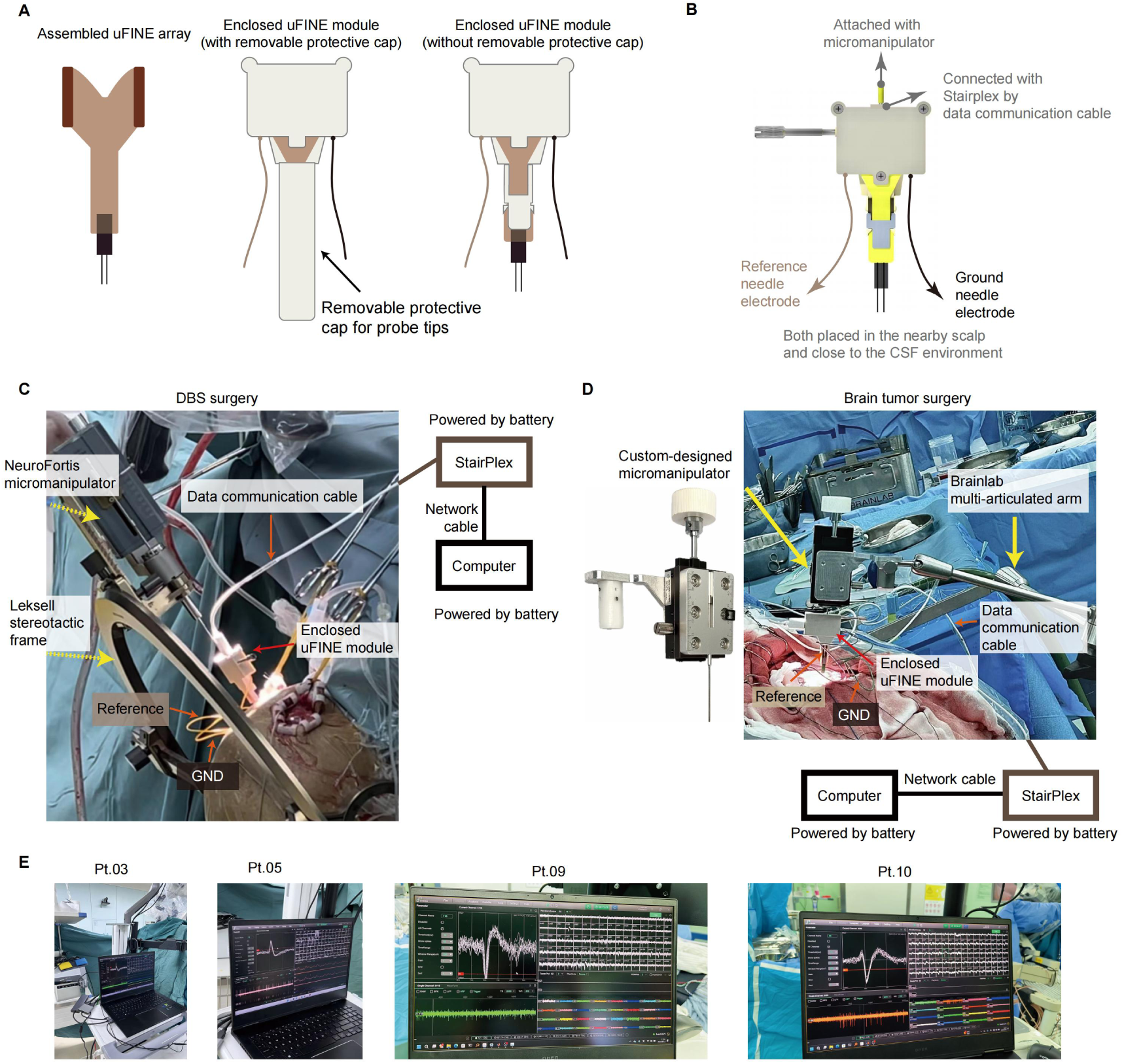
Array implantation and recording setup in different intraoperative settings. (A) Schematic diagram showing assembled uFINE array and enclosed uFINE module. Before implantation, the uFINE array tip was protected by a removable protective cap. (B) Schematic diagram showing the connection setup of the enclosed uFINE module. (C) Implantation setup in DBS surgery. The enclosed uFINE module were attached to the NeuroFortis micromanipulator, which was mounted on a stereotactic frame for precise positioning during the procedure. (D) Implantation setup in tumor resection surgery. (E) Photos showing the spike waveforms displayed on the StairPlex user interface during intraoperative recordings in Pt.03, Pt.05, Pt.09, and Pt.10.

To accommodate various neurosurgical procedures, we developed different implantation setups for the uFINE array (Figure 1C and Figure S2C-D). For the cases of DBS implantation, we designed an adaptor that could be mounted onto the Leksell stereotactic frame and coupled with commercial micromanipulators for precise array insertion (Figure 1C and Figure S2C). This setup allowed for insertion through the burr hole, minimizing unnecessary adjustments of the stereotactic frame and preventing additional tissue damage. During surgery, the pre-assembled and sterilized uFINE array and headstage were securely attached to the adaptor by the surgeon. In brain tumor resection surgeries, a custom-designed micromanipulator, mounted on a commercial multi-articulated arm, was used for array insertion (Figure S2D). In all patients, a small superficial incision in the pia was done at the insertion site. During implantation, the uFINE array was guided by coupled tungsten shuttle wires (75 µm in diameter) and inserted to a depth of 5-6 mm. Once the target depth was reached, the shuttle wires were withdrawn, leaving only the flexible array shanks embedded in brain tissue. Therefore, although the back end of the uFINE array was fixed to the holding apparatus, the flexible shanks could follow brain pulsations, which effectively mitigated relative motion between the brain tissue and the recording sites (Figure 1D).

The uFINE array used in intraoperative recordings contained 128 recording sites of 50 μm circle in diameter with 65-µm spacing distributed over two shanks (Figure S1A). Each shank measured 20 mm in length, 80 to 245 µm in width with a tapered structure, and 2 µm in thickness, with a typical spacing of 1 mm between the shanks. The 64 recording sites on each shank covered a 4-mm distance, enabling dense sampling of neural activity at single-cell resolution across cortical depths in the human brain. Compared to penetrating neural electrodes that have been used in human (e.g., microwire bundles and Neuropixels probes), the uFINE array features micron-level dimensions, superior flexibility, and mechanical robustness (Figure 1E-H), making it well-suited for intraoperative use. The array shank offered tensile strengths of up to 831.78 ± 67.87 MPa (mean ± std, n = 5, Figure 1H), which is critical for minimizing the risk of breakage during implantation and ensuring patient safety. The impedance of the uFINE recording sites at 1000 Hz was 52.99 ± 0.49 kΩ (mean ± std, n = 6, Figure 1I).

### Single-unit recording of neuronal populations across cortical depths

The linear arrangement of uFINE recording sites allowed for sampling of neural activity across cortical depths in the human brain. As shown in Figure 2A-B, recordings from Pt.05 exemplify both low raw signals and high-pass filtered (250 Hz) spiking activity captured by the recording sites distributed along the array shanks. Multiple single units with distinctive waveforms were isolated using Kilosort 4^36^ followed by manual curation (Methods). A substantial fraction of single units was detected by more than one recording site, facilitating accurate spike sorting^37^ (Figure 2C).

**Figure 2.**
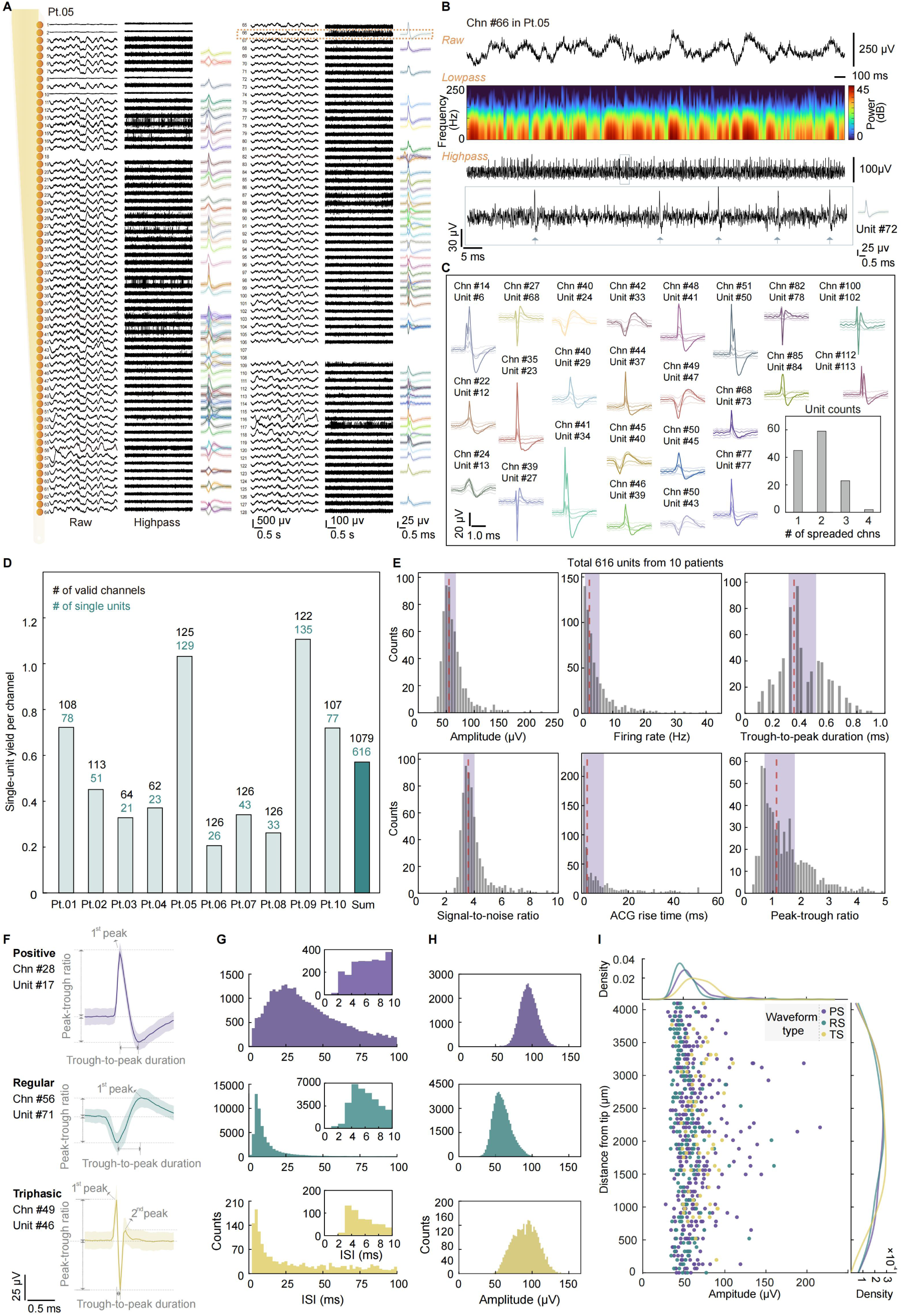
Single-unit recordings across depths in human cortex. (A) Examples of 2-s recording segments from Pt.05, showing the raw signals and spiking activity (highpass-filtered, >250 Hz) recorded by two shanks of the same uFINE array. The waveforms of individual single units are located near the channel with their largest amplitudes. The average waveform and std (shaded areas) were calculated from 500 spikes for each single unit.The leftmost schematic diagram shows the distribution of recording sites. (B) Raw signals, the spectrogram of LFP signals (lowpass-filtered, <250 Hz), and spiking activity from example Chn #66, corresponding to the orange dashed box in (A). Zoomed-in view shows the waveform of individual spikes (marked by asterisks) of Unit #72. Chn, channel. (C) Waveform spread of example units from Pt.05. Waveforms from 5 adjacent channels are displayed for each unit. Inset, the number of units detected by different numbers of channels. (D) Single-unit yields of each patient. The numbers above each column indicate the number of valid channels and the number of single units. The rightmost column represents the summation from all patients. (E) Quality metrics of 616 single units pooled from the 10 patients. Red dashed lines indicate the median, and the purple shaded areas represent the interquartile range for each metric. (F) Waveforms of representative units from the three classes: positive spiking (top, purple), regular spiking (middle, green), and triphasic spiking (bottom, yellow). The average waveform and std (shaded areas) were calculated from 500 spikes. (G-H) Distributions of the inter-spike interval (G) and amplitude (H) of the example single units shown in (F), color coded as in (F). Insets, zoomed-in views showing the distribution between 0-10 ms. (I) Relationship between the mean amplitude (x axis) and the distance from the array tip (y axis, calculated by multiplying the channel number by site spacing) of each single unit. Kernel density estimation (KDE) plots for both metrics are shown along the x and y axes. Unit classes are color coded as in (F).

**Figure S3.**
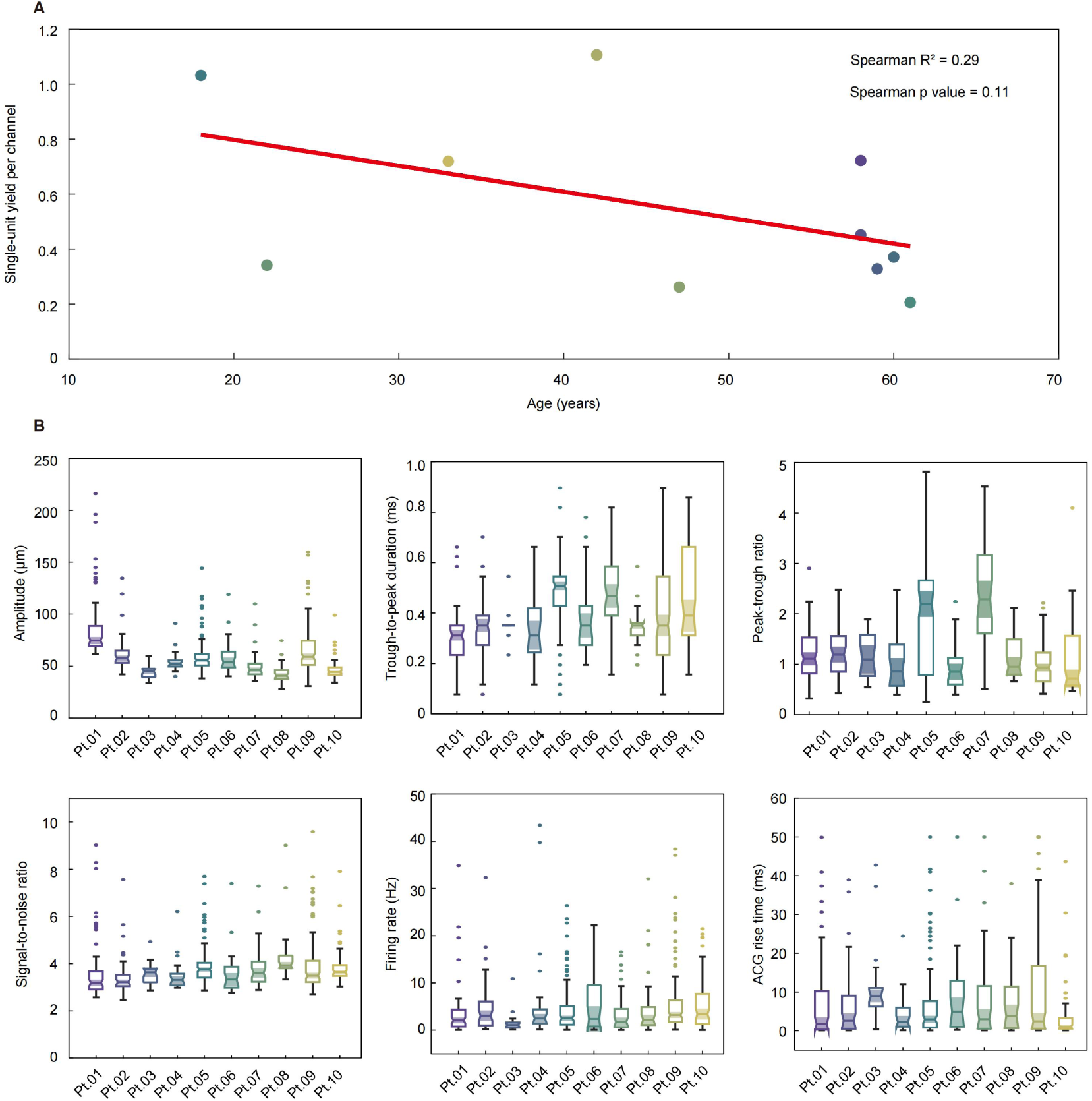
Single-unit yield per channel and quality metrics in individual patients. (A) The single-unit yield per channel shows a decreasing trend as the patient age increases (p = 0.11, R² = 0.29, Spearman correlation analysis). Each dot represents the result from one patient. (B) Quality metrics quantified for individual patients. Boxplots indicate the median, the interquartile range, and the range of values. Outliers are shown as individual data points outside the whiskers. Shaded areas indicate the notch.

In total, 616 single units were isolated from 1079 valid channels (impedance < 4 MΩ) across 10 patients, yielding an average of 0.55 ± 0.32 single units per channel (Figure 2D). We also observed considerable inter-patient variability in single-unit yield per channel, with a trend of decreased yield with aging (p = 0.11, R² = 0.29, Spearman correlation analysis, Figure S3A), consistent with previously reported age-related declines in neuronal density^38^. The quality of single units was evaluated by a set of metrics (Figure 2E; see Figure S3B for individual patient metrics). The median and interquartile range of each quality metric were as follows: amplitude: 54.85, 46.45 to 66.90 μV; firing rate: 2.50, 1.08 to 5.74 Hz; signal-to-noise ratio (SNR): 3.57, 3.22 to 4.00; autocorrelogram (ACG) rise time: 2.39, 0.63 to 9.52 ms; trough-to-peak duration: 0.35, 0.31 to 0.51 ms; and peak-trough ratio: 1.13, 0.70 to 1.78. These values are comparable to those reported in previous primate studies^19^.

Although spike waveforms depend on both neuronal type and the location of the recording site relative to the soma, many neuroelectrophysiological studies still infer putative neuronal types from waveform characteristics to better understand neural mechanisms^20,39^. Based on the polarity and peak-trough ratio of the largest waveform of each single unit (Methods), we classified these units into three categories (Figure 2F-H): positive spikes (PS, n = 372 single units), regular spikes (RS, n = 178), and triphasic spikes (TS, n = 66), which putatively correspond to axonal action potentials, excitatory neurons, and inhibitory neurons, respectively. The spatial distribution of these three classes across penetration depths showed no significant difference (p = 0.76, Kruskal-Wallis test). TS units exhibited slightly larger amplitudes (68.35 ± 15.71 μV, mean ± std) compared to RS units (62.35 ± 24.63 μV, mean ± std, p = 0.01, Welch t test) and PS units (53.61 ± 18.06 μV, mean ± std, p = 5.37 × 10^−9^, Welch t test) (Figure 2I).

Taken together, these results demonstrate that the uFINE array is capable of achieving high-quality recordings of single units at scales in the human brain.

### Stable intraoperative recording of single-unit activity

A major challenge for intraoperative recordings using rigid electrodes is the substantial human brain pulsations caused by respiratory and cardiac cycles^19,20^. These rapid brain movements, usually fluctuating around respiratory frequency, pose difficulties for automatic spike-sorting approaches^40,41^ and have been shown to negatively correlate with single-unit yield^19^. However, the flexibility of the uFINE array could address this issue by allowing its shanks to follow brain movements, effectively minimizing relative motion between the brain tissue and recording sites, and thereby enabling stable single-unit recordings (Figure 1D).

As shown in Figure 3A, the spike position remained stable over time in a representative recording segment, with no discernible cycles reflecting brain pulsations. To further estimate the effects of rapid brain movements on recording stability, we identified a subset of single units showing high firing rates (>0.5 Hz) and large amplitudes (>70 µV) as spatial markers (n = 112 single units from 9 patients) to observe their spike position drift (a segment from Pt.10 is shown in Figure 3A). The average standard deviation of spike position of these units was 18.88 ± 7.69 µm (mean ± std, Figure 3B), indicating minimal displacement due to rapid brain movements. In addition, for all single units, we also estimated the distance between spikes of each single unit and its nearest neighbouring site (the recording site with the largest waveform for this unit identified from spike sorting) over the course of recordings (Methods). The majority of all single units (60.84% of 616 single units) remained within 32.5 µm of their nearest neighbouring site (Figure 3C), further supporting the minimized relative motion between the recording sites and the brain tissue. In a subset of patients (n = 6) with concurrent video recordings of the brain surface during array insertion, we further confirmed that the amplitude of rapid brain movements (Figure S4A) had no significant impact on either the single-unit yield (R² = 0.23, p = 0.34) or the single-unit yield per channel (R² = 0.28, p = 0.28) (Figure S4B).

**Figure 3.**
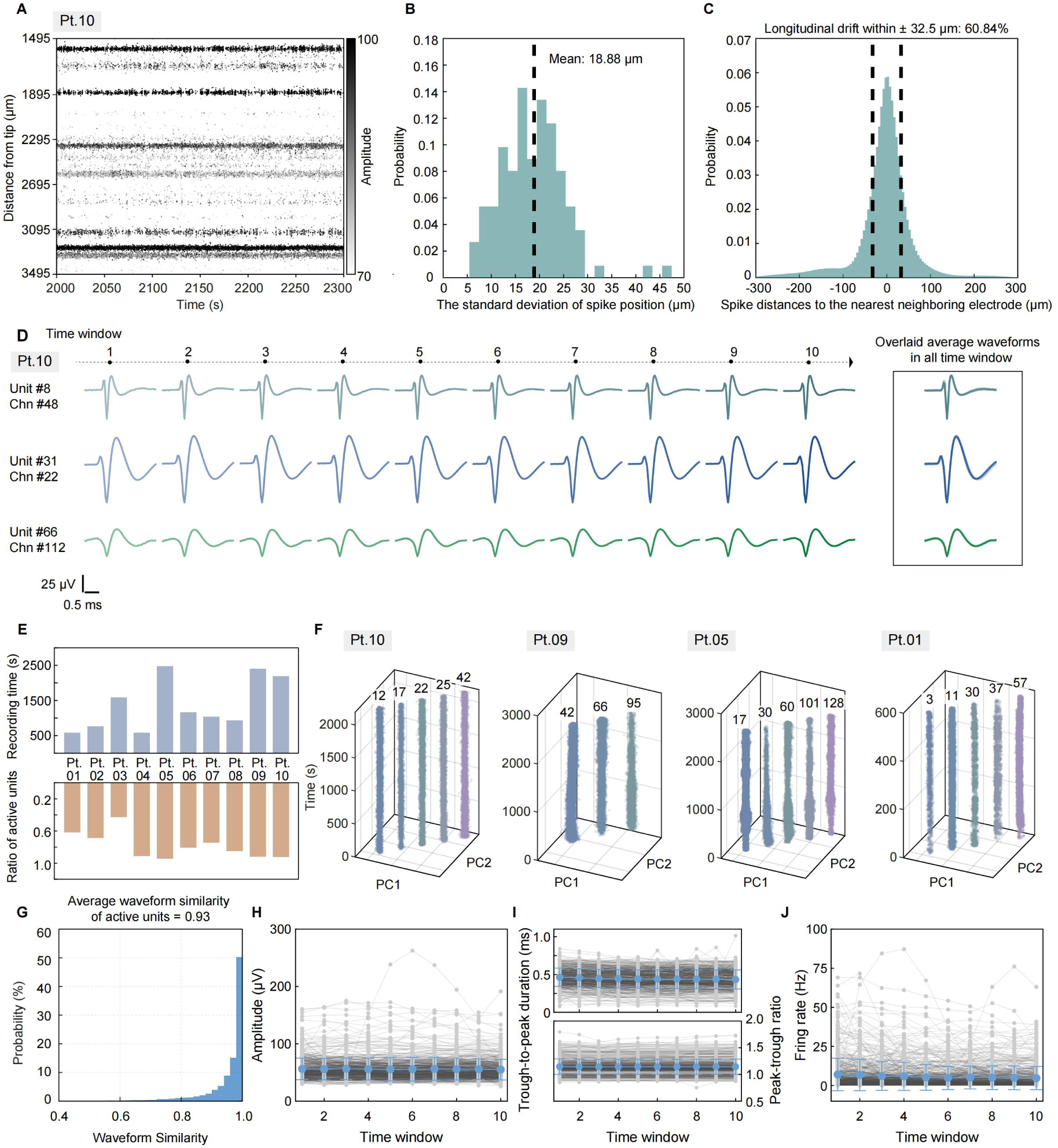
Stable intraoperative single-unit recording. (A) Example recording segment from Pt.10. showing spike positions over time. Each dot represents a detected spike. The array tip corresponds to the 0 mm. (B) Histogram showing standard deviation of spike positions within each unit (n = 112 units as spatial markers from 9 patients). The black dashed line marks the mean value. (C) Histogram showing spike positions relative to the nearest neighboring site of individual units throughout the recording (n = 616 single units). Black dashed lines indicate the range of ± 32.5 µm (half of the site spacing). (D) Left: Average waveforms of three representative active units in Pt.10 throughout the recording. Each waveform represents the average of 500 spikes in the corresponding time window. Right: Average waveforms of the same example units overlaid across time windows. (E) Recording duration (top) and ratio of active units (bottom) of each patient. (F) Time evolution of example single-unit waveforms (from Pt.10, Pt.09, Pt.05, and Pt.01.) projected in the 2D PC space. The x and y axes denote dimensions corresponding to the first and the second PC, respectively, and the z axis denotes the recording time. (G) Waveform similarity of active units pooled across all recording sessions (n = 511 active units from 10 patients). (H)-(J) Time evolution of amplitude (H), trough-to-peak duration (I, top), peak-trough ratio (I, bottom), and firing rate (J) of active units (n = 511). Blue dots represent the mean value across active units. Gray dots represent the value of individual active units. Error bars, std.

**Figure S4.**
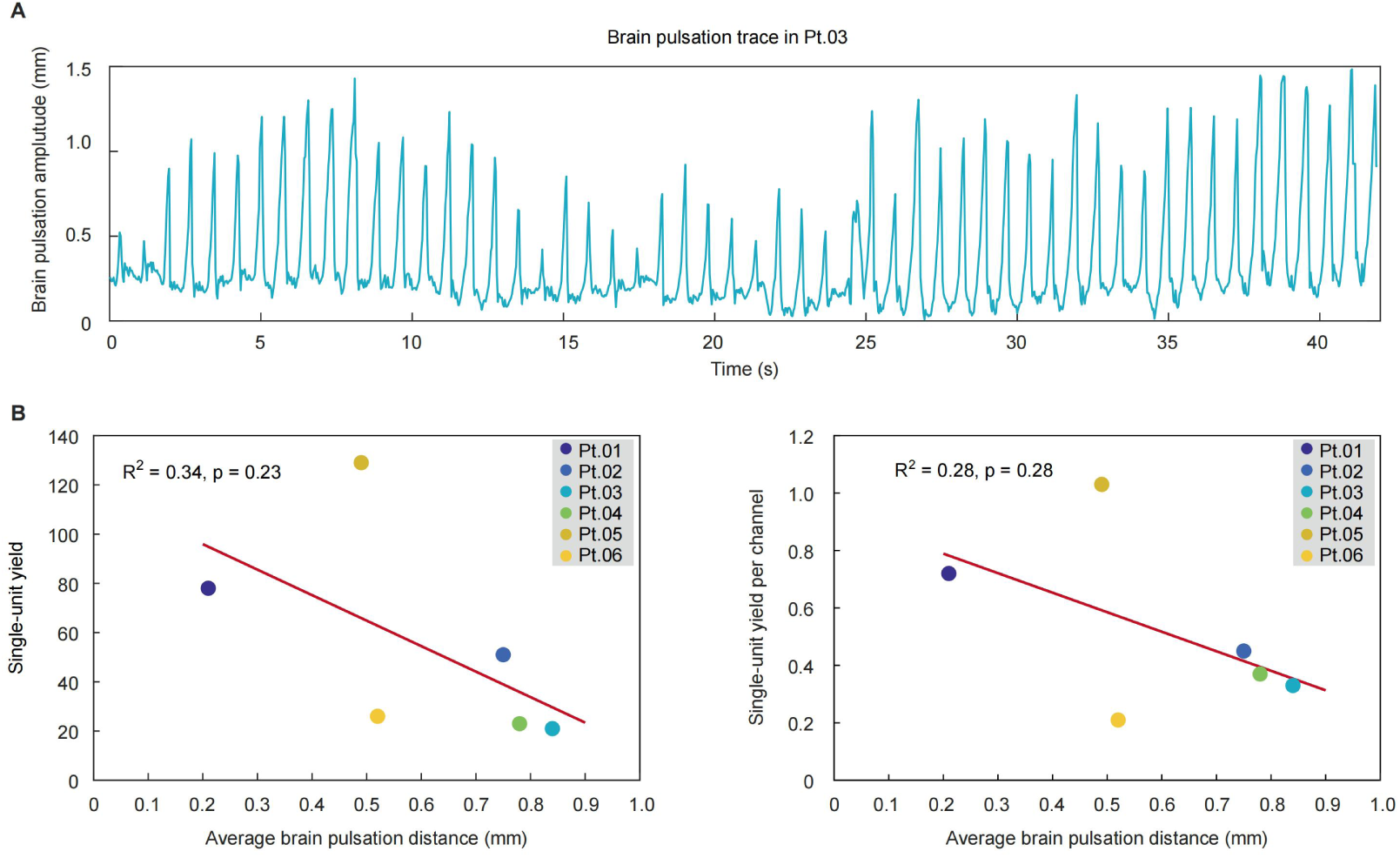
The single-yield yield was not affected by brain pulsations. (A) A representative trace of brain pulsations extracted from video recordings from Pt.03. (B) Neither the single-unit yield (left, R² = 0.34, p = 0.23, Pearson correlation analysis) nor the single-unit yield per channel (right, R² = 0.28, p = 0.28, Pearson correlation analysis) was negatively correlated with the amplitude of brain pulsations. Dots are color coded and represent the results from individual patients with video recordings.

The mitigated relative motion between the brain tissue and the recording sites should also help preserve neuronal health, thereby facilitating stable capturing of neuronal activity over longer timescales (e.g., minutes). As shown in Figure 3D, the waveforms of individual single units maintained high similarity over the course of a 36.5-minute recording from Pt.10. To evaluate the recording stability throughout the intraoperative recording period, we divided each recording into 10 time windows and focused on single units with sufficient spike events (termed “active unit”; firing rate > 0.1 Hz and at least one spike in each time window, Methods). On average, 78.30 ± 16.69 % of single units across 10 patients met this criterion (totaling 511 units, Figure 3E and Table S1). The waveforms of each active unit occupied nearly consistent positions in a low-dimensional PC space, indicating that waveform shape of individual units remained stable over the course of recording (Figure 3F). Throughout the recording, active units exhibited high waveform similarity within each unit (0.93 ± 0.14, mean ± std, Figure 3G), along with excellent stability in spike amplitudes, trough-to-peak durations, peak-trough ratios, and firing rates (Figure 3H-J; see statistics in Table S2). Taken together, these results demonstrate the capability of the uFINE array to achieve stable single-unit recordings over both fast and prolonged timescales in the intraoperative environment.

### Stimulus and response tuning at single-cell level in human dlPFC

The capability to record from populations of neurons with single-cell resolution provides a valuable opportunity to investigate sensory, motor, and cognitive processes in human brain. In patients undergoing DBS lead implantation, the uFINE array was implanted in the dlPFC, an area has extensive connections with sensory and motor cortices and has been implicated in working memory, cognitive flexibility, and planning^42–45^. We began by characterizing the activity modulation of individual neurons in human dlPFC using a simple “Word-Repeating” task (Figure 4A, 4B). During each trial, the patient (Pt.03, Pt.04 and Pt.08) listened to a word (also displayed on a screen) for 0.5-1.0 s and was then required to orally repeat the word after a 0.6-s delay. On average, each patient completed 79.67 ± 21.01 (mean ± std) trials. We found that the activity of some single units was significantly modulated during the stimulus and response period (Figure 4C-N and Methods), with a slightly higher fraction of units modulated during the response period than the stimulus period (39.10 ± 24.68% vs 22.83 ± 12.64%, mean ± std, Figure 4F, 4J and 4N). Overall, more dlPFC single units exhibited suppression rather than activation (19.65 ± 11.64% vs 3.17 ± 5.50% for the stimulus period, 34.00 ± 28.49% vs 5.10 ± 5.27% for the response period, mean ± std, Figure 4F, 4J and 4N). Our intraoperative recordings highlighted heterogeneous single-unit responses during task performance, showing the potential to facilitate detailed explorations of the neural mechanisms underlying various human cognitive functions at the cellular level.

**Figure 4.**
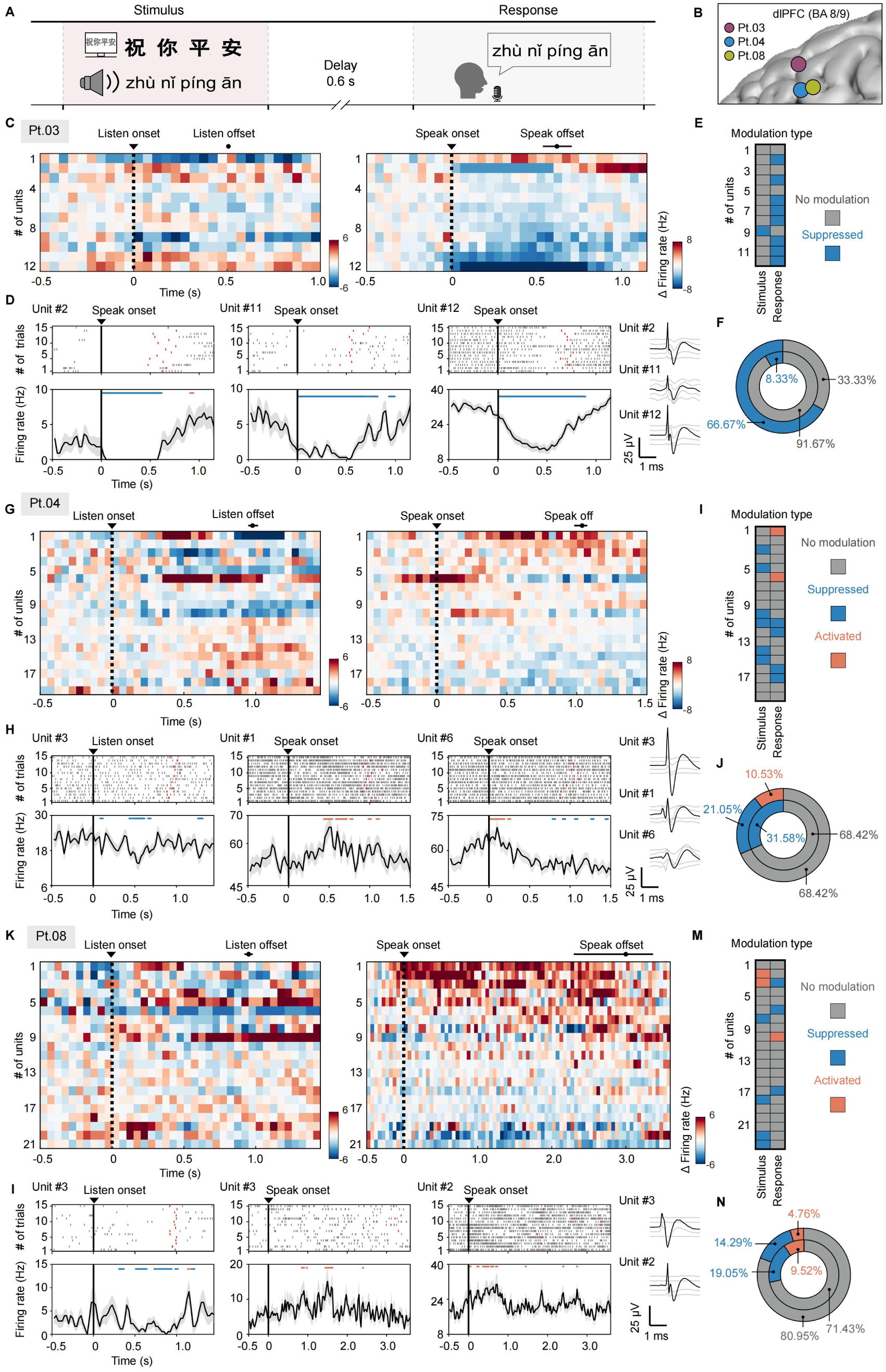
Stimulus and response tuning at single-cell level in human dlPFC. (A) Schematic diagram showing the structure of the “Word-repeating” task. (B) The implantation site of the uFINE array in the three patients involved in this task, all located in the dlPFC. BA, Broadmann area. (C) Trial-averaged activity changes of individual units aligned to “listen” or “speak” onset. Black dots and horizontal lines indicate the average offset timing and 25-75% range, respectively. (D) Task-related activity modulation of example single units marked by asterisks in (C). Top: raster plots (spikes). Red ticks indicate the offset timing in each trial. Bottom: trial-averaged firing rates. Horizontal lines indicate time bins when the unit was significantly activated (red) or suppressed (blue) (Wilcoxon signed-rank test, p < 0.05). The rightmost panels are waveform spreads of example units. (E) Types of activity modulation of individual units (ordered as in (C)) during the stimulus and the response period. (F) Fraction of units with significant activity changes during the stimulus (outer circle) and the response period (inner circle). The color codes are the same as those in (E). (G-J) Same as (C)-(F) but for Pt.04. (K-N) Same as (C)-(F) but for Pt.08.

## DISCUSSION

In this study, we introduced a method that leverages ultra-flexible microelectrodes, specifically the uFINE array, for acute single-unit recordings in the human brain during intraoperative procedures. Our findings demonstrate that the uFINE array enables stable, high-quality recordings across cortical depths, even in the presence of rapid brain movements. This recording capability allowed us to characterize the heterogeneous responses of individual dlPFC neurons in a “Word-Repeating task”. These results highlight the potential of the uFINE array as a powerful tool for intraoperative neurophysiological investigations.

### Advantages of the uFINE array for intraoperative use

The single-unit yield of the uFINE array is comparable to that of microelectrodes with higher channel counts (e.g., Neuropixels probe)^19–23^. Notably, the uFINE array shows a higher single-unit yield per channel, indicating that reliable data collection can be achieved with fewer channels. This recording efficiency stems from the combination of superior flexibility and reduced invasiveness.

One of the primary advantages of the uFINE array over other recording technologies is its exceptional tolerance to brain pulsations, which often lead to significant signal fluctuations. Rigid electrodes typically require intraoperative adjustments, such as lowering PEEP (positive end-expiratory pressure) to mitigate pulsation effects, and offline alignment algorithms to correct movement artifacts^19,20,24^. These methods, however, may not fully eliminate the negative impact of brain pulsations on single-unit detection. In contrast, previous studies have shown that ultra-flexible microelectrodes exhibit bending stiffness comparable to single-cell traction forces^26^. This allows the uFINE array to follow brain movements in real-time without the need for additional adjustments, either intraoperatively or during post-processing, significantly enhancing recording stability. Thus, although the current design of the uFINE array does not offer the same electrode density as Neuropixels probes, it still enables stable tracking and isolation of individual units. Additionally, its ability to adapt to brain movements reduces friction between the electrodes and the brain tissue, minimizing tissue damage and preserving neuronal health during prolonged recordings.

In addition to flexibility, the uFINE array minimizes tissue damage, which is critical for acquiring high-quality signals in acute recordings. The excellent mechanical robustness of the uFINE array allows it to maintain structural integrity while keeping small dimensions even when used in the human brain. The insertion foot print of the uFINE array, with and without shuttle wires, is 4.91 × 10^-9^ m^2^ and 4.90 × 10^−10^ m^2^, respectively. The cross-sectional area of the uFINE array shank is 14.29 times as small as that of the Neuropixels 1.0-S probe^20,24^, ∼100 times smaller than that of single tungsten microelectrode arrays^46^; and ∼2000 times smaller than that of SEEG-coupled microwire bundles^8–11^. This reduction in cross-sectional area results in a more tissue-friendly implantation, allowing for faster recovery of neural activity after insertion.

### Limitations and future directions

Despite the advantages, the overall recording scale in this study was still limited. Two out of ten of the recordings yielded more than 100 single units, partly because each uFINE array currently contains only 128 recording sites. Future iterations of the uFINE array, with more recording sites and optimized electrode configurations, will help expand the scale of recording. In addition, the tungsten shuttle wires created an insertion track much larger than the uFINE array shank. Using finer tungsten wires could further minimize insertion damage and reduce the time required to obtain stable neural signals, which is particularly critical in time-sensitive intraoperative recordings. Moreover, the long-term performance of the uFINE array, particularly its ability to maintain stable recordings over extended periods (e.g., months to years), requires further investigation.

### Prospects of future applications

Once these limitations are overcome, we anticipate that the uFINE array will have broad applications in both neuroscience research and clinical practice. One of its key advantages is the ability to record from many neurons across different cortical layers. The capacity to capture high-resolution neuronal data opens up new avenues for exploring human brain activity in both healthy and diseased states. In our study, we revealed the heterogeneous responses of individual neurons in the dlPFC to sensory and motor events. Future studies with larger-scale single-unit recordings will enable more comprehensive investigations of complex cognitive functions in the human brain.

Beyond basic neuroscience, the uFINE array also has significant potential in clinical applications. Its ability to record from individual neurons in brain regions implicated in brain disorders could offer valuable insights into neurodegenerative diseases, psychiatric disorders, and brain injury. Moreover, the uFINE array could play a pivotal role in future brain-machine interfaces (BMIs) by providing stable and reliable neural signals for controlling prosthetics or other assistive devices. Its capacity for large-scale, long-term recordings with minimized tissue damage positions it as a crucial tool for advancing BMI technologies.

In summary, the uFINE array represents a significant advancement in the field of intraoperative neurophysiology, offering a stable, high-yield solution for recording single-unit activity with minimized tissue damage in the human brain. As the technology evolves, the uFINE array holds great promise for advancing both basic neuroscience and clinical applications.

## RESOURCE AVAILABILITY

All data are available in the main text or the supplementary materials.

### Lead contact

Further information and requests for resources and reagents should be directed to and will be fulfilled by the lead contact, Xue Li.

### Materials availability

This study did not generate unique reagents.

### Data and code availability

- The raw datasets supporting the current study have not been deposited in a public repository because they contain personally identifiable patient information, but they are available in an anonymized form from the lead contact upon reasonable request.
- Any additional information required to reanalyze the data reported in this paper is available from the lead contact upon request.

## ACKNOWLEDGMENTS

We thank Xiaocheng Li and Nanofabrication Facility for Advanced Brain Science at CEBSIT for supporting electrode fabrication; Xingyu Liu for discussing paradigm design and optimization; Bingbing Li for refining visual representations; Xing Shao for designing the implantable casing and the micro-drive system; Guangyao Zhang for “Word-repeating” task coding; Zexing Zhao for his work in data acquisition; Zhuo Chen for processing and analyzing the collected data; Zeyu Wang for assisting with electrical impedance spectroscopy (EIS) characterization; Shun Bai, Kang Meng and all members in our lab for recommendations on electrophysiological recording techniques; Shanghai Stairmed Technology Co.,Ltd. for supporting electrode fabrication, hardware & software support and medical communication; and the clinical teams for their invaluable collaboration and support. We are deeply grateful to the patients who participated in this study, whose contributions were essential for this research. Z.Z. is supported by Shanghai Municipal Science and Technology Major Project grant 2018SHZDZX05, Shanghai Municipal Science and Technology Major Project grant 2021SHZDZX and National Science and Technology Innovation 2030 Major Program grant 2022ZD0210300. X.L. is supported by National Science and Technology Innovation 2030 Major Program grant 2021ZD0202202 and Shanghai Municipal Science and Technology Major Project grant 2021SHZDZX.

## AUTHOR CONTRIBUTIONS

Conceptualization, X.L, Z.Z; Methodology, X.L, Z.Z, S.W, Z.Y, X.L, J.L; Investigation, S.W, X.L, Z.Y, X.L, Y.Q, J.L, Q.D, C.K, G.C, B.C, Z.Z; Data curation, S.W, C.K; Visualization, C.K, S.W; Writing— original draft, C.R, S.W, C.K; Writing—review & editing, Z.Z, S.W, J.L, X.L, X.L, X.J; Supervision, X.L, Z.Z, C.R; Project administration, X.L, Z.Z, C.R, S.W; Funding acquisition, X.L, Z.Z.

## DECLARATION OF GENERATIVE AI AND AI-ASSISTED TECHNOLOGIES

During the preparation of this work, the authors used GPT-4o in order to assist code development, as well as to refine and proofread textual content. After using this tool, the authors reviewed and edited the content as needed and take full responsibility for the content of the publication.

## TABLES AND TEXT BOXES

**Table S1.**
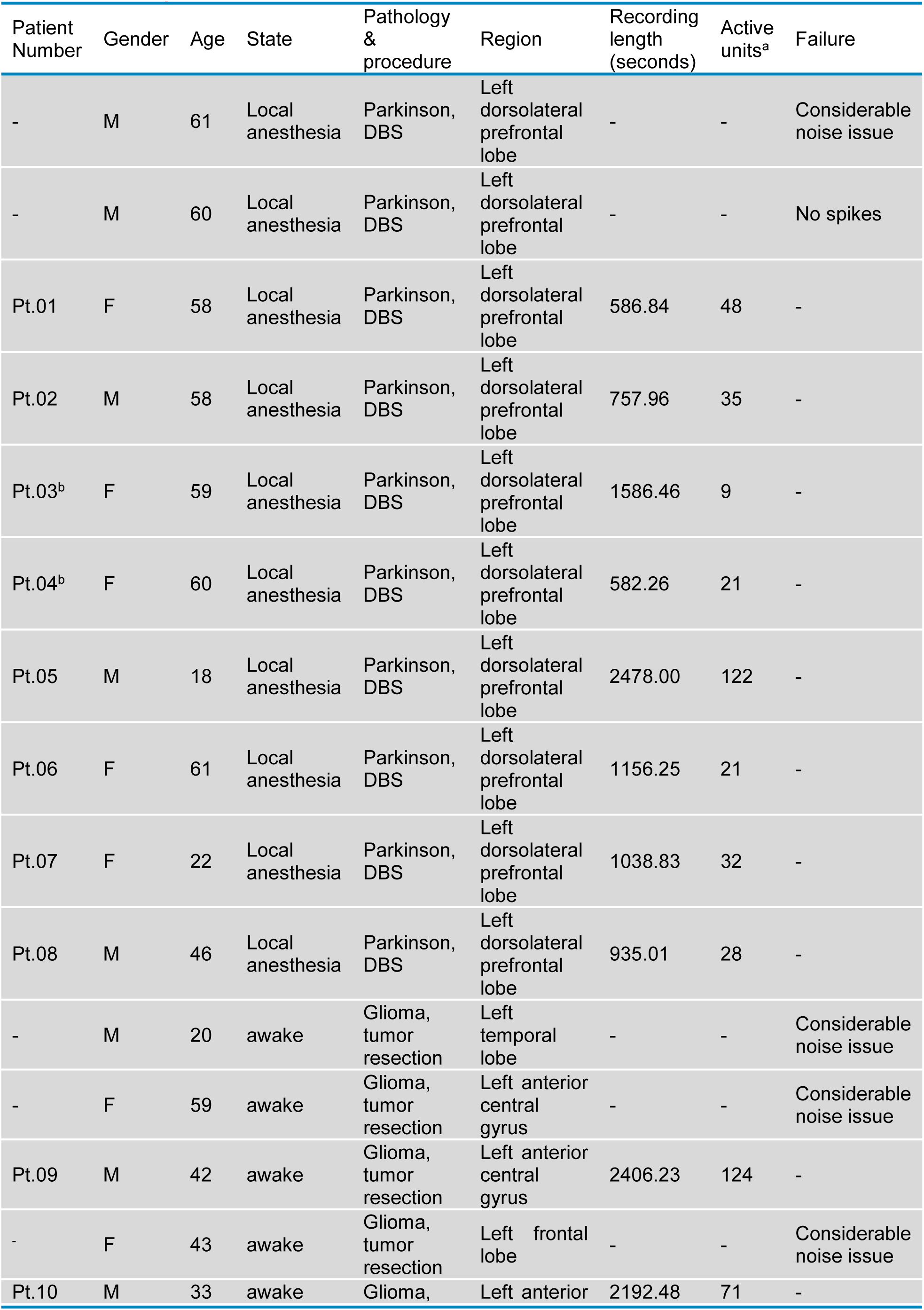
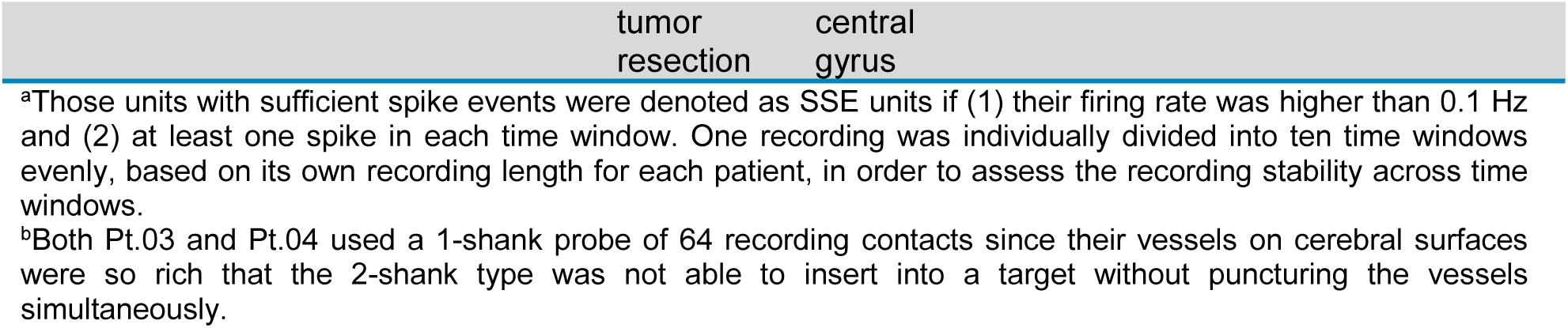
Participant information.

**Table S2.** Statistics of active unit characteristics across time windows for each patient^a^.

| Patient<br>Number | Amplitude |  | Firing rate |  | Trough-to-peak<br>duration |  | Peak-to-trough ratio |  |
| --- | --- | --- | --- | --- | --- | --- | --- | --- |
|  | p-value | Significant | p-value | Significant | p-value | Significant | p-value | Significant |
| Pt.01 | 0.74 | N<sup>b</sup> | 0.86 | N | 0.01 | Y | 1.00 | N |
| Pt.02 | 0.43 | N | 0.99 | N | 0.89 | N | 1.00 | N |
| Pt.03 | 0.30 | N | 0.02 | Y | 0.00 | Y | 1.00 | N |
| Pt.04 | 0.99 | N | 0.97 | N | 0.99 | N | 1.00 | N |
| Pt.05 | 0.07 | N | 0.01 | Y | 0.74 | N | 1.00 | N |
| Pt.06 | 0.98 | N | 1.00 | N | 0.99 | N | 1.00 | N |
| Pt.07 | 0.10 | N | 0.65 | N | 0.89 | N | 0.99 | N |
| Pt.08 | 0.91 | N | 1.00 | N | 0.93 | N | 1.00 | N |
| Pt.09 | 0.97 | N | 0.00 | Y | 0.07 | N | 1.00 | N |
| Pt.10 | 0.58 | N | 1.00 | N | 0.56 | N | 1.00 | N |

<sup>a</sup>ANOVA performed to compare metrics across time windows (Methods).No significance if p>0.05.
<sup>b</sup>N = No significant difference; Y = Significant difference.

## STAR★METHODS

### KEY RESOURCES TABLE

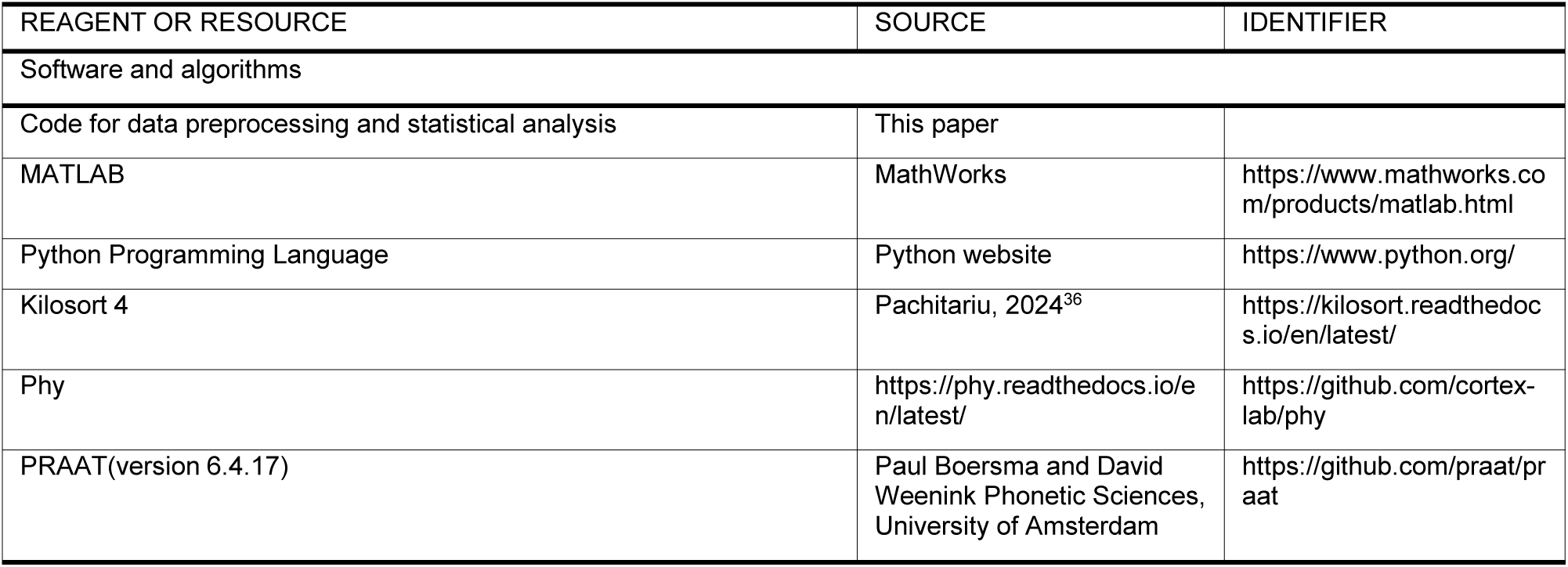

### EXPERIMENTAL MODEL AND STUDY PARTICIPANT DETAILS

The protocols were approved by the Medical Ethics Committee of the First Affiliated Hospital of the Air Force Medical University (KY20232366-F-1) and the Huashan Hospital Institutional Review Board of Fudan University (HIRB, KY2024-103).

Fifteen participants were included in the study across two hospitals (Table S1), all of whom were already scheduled for intraoperative neurophysiological procedures during either DBS implantation surgery under local anesthesia or tumor resection with awake functional mapping as part of normal clinical routine. There was no additional risk of participants.

All participants provided written informed consent and were free to withdraw from the study at any time, including during surgery, without any impact on their clinical care.

## METHOD DETAILS

### Design, fabrication and assembly of uFINE electrode array

The uFINE array used in this study consisted of 128 channels arranged in two shanks with shank spacing of 1000 μm. The shank thickness was 2 μm, and the length of a single shank was 20 mm. Each shank featured 64 recording sites, vertically aligned along its front end, as the recording site segment. The total length of this segment was approximately 4.15 mm, with each circular site having a diameter of 50 μm and a center-to-center distance of 65 μm. The recording site segment had a tapered structure, with a maximum width of 245 μm at the implantable portion and a width of 80 μm at the tip. Besides the recording site segment, the rest of the shank maintained sufficient redundancy to ensured that it remains in a slack state during brain movement, preventing the portion of the shank inside of the brain from being pulled out.

The uFINE was fabricated using standard planar microfabrication techniques (see Figure S1B). Detailed descriptions of the fabrication and assembly methods were provided in previous publications^29^. This study implemented several improvements based on prior work, achieving higher uFINE array tensile strength through fabrication and utilizing medical-grade materials throughout the assembly process. The specific enhancements were as follows:

A. To further cure the polyimide (PI) and enhance the structural integrity of the electrodes, the PI hard baking temperature was increased from 300°C to 320°C, and the curing time was extended from 30 minutes to 2 hours.
B. The diameter of the tungsten shuttle wires used for auxiliary electrode implantation was increased from 50 μm to 75 μm.
C. A bio-compatible surgical adhesive (Kwik-sil, World Precision Instrument Co.) was used to secure the shuttle wires to the silicon substrate. (In contrast to previous methods, this project did not require a needle withdrawal design during surgery.)
D. In this study, the “assembled uFINE array” (Figure S2A) referred to the state in which the uFINE array was anchored to tungsten shuttle wires and soldered to a flexible printed circuit (FPC) for connection between headstage and uFINE array.
E. In this project, after the tungsten shuttle wires were retracted from the brain tissue, there was no need to remove them from the silicon substrate that held the tungsten shuttle wires; instead, they were left suspended 3-5 mm above the brain surface to ensure that they did not scratch the brain surface.
F. A waterproof protective case was specifically designed for complete sterilization, encapsulating the assembled uFINE array and headstage (128 channels, Shanghai Stairmed Technology Co.) together, to form an enclosed uFINE module (Figure 1B, Figure S2A). This case was made from a common medical material, polyether ether ketone (PEEK), with reference and ground wires made from needle electrodes (Xian Fude Co.) connected to the headstage. The case consisted of four parts, including a removable protective cap to prevent potential damage to the uFINE array tip before implantation (see Figure S2A). The top of the case featured a port for connecting power and data lines. Prior to implantation, the enclosed uFINE module could be connected to a NeuroFortis (AlphaOmega Co.) micromanipulator fixed to a stereotactic frame (Leksell Co.) or to a custom-designed micromanipulator mounted on a commercial multi-articulated arm (Brainlab Co.).
G. We also prepared a 1-shank array to avoid complications related to vascular avoidance during the implantation of the 2-shank array. The assembly process for the 1-shank array was similar to that of the 2-shank array; however, the 1-shank array utilized only one tungsten shuttle wire to secure one of the uFINE shanks, resulting in a total of 64 recording sites. Pt.03 and Pt.04 in this study were implanted with the 1-shank array.

### In vitro characterization of the uFINE array

#### (1) Mechanical

Tensile test of the uFINE array was conducted using an electronic balance (BSM-220.3S, Shanghai ZhuoJing Co.), a 50g weight, and a Digital Stereotaxic Instrument (68025, RWD).

One shank of the uFINE array was released and cut from the wafer. Both ends of the shank were bonded separately to the weight and the holder of the stereotactic device using adhesive. The middle section of the shank was maintained at a length of 1.4±0.1 cm in a relaxed state. The weight was placed on the balance.

The stereotactic device was gradually raised with constant speed monitored by digital display module, stretching the electrode shank and causing slight displacement of the weight, which was reflected in changes to the balance readings. These readings were recorded until the electrode shank broke. The maximum force the shank could withstand was calculated based on the change in balance readings and the electrode’s cross-sectional area, yielding the maximum tensile stress the electrode could endure. This experiment was repeated five times. Results are shown in Figure 1H.

#### (2) Electrochemical Impedance Spectroscopy

To assess the electrode’s ability to maintain good recording performance across a broad frequency range (i.e., capable of recording neural activity across most frequency bands), a Gamry Reference 620 potentiostat (Gamry Instruments) was used to measure the impedance and phase angle of the electrode over a frequency range from 10 Hz to 10^4^ Hz. The results are shown in Figure 1I.

### Preoperative preparation

#### (1) Sterilization and scenario setup

The items requiring sterilization before surgery included the enclosed uFINE module (containing the assembled uFINE array, FPC, and headstage), data communication cable (Shanghai Stairmed Technology Co., Ltd.), micromanipulators, and a multi-articulated arm (for tumor resection surgery). These items were sterilized in the hospital using ethylene oxide (EO) sterilization following standard protocols.

As shown in Figure S2B-D, other items that needed to be prepared included the StairPlex recording system (Shanghai Stairmed Technology Co., Ltd.), laptop (Windows 11, IE9), battery, and equipment required for behavioral tasks (speakers, microphone, and NI Data Acquisition Card (National Instruments, USB-6009)). These components were transported into the operating room using a custom-assembled cart and connected prior to the experiment. Specifically, the StairPlex, speakers, microphone, and NI card were connected to the computer.

#### (2) Denoise

Denoising of intraoperative acute recording included two key operations after careful noise signal mitigation tests:

1. Set the ground of headstage connected into the scalp.
2. Eliminate the power line interference from recording devices.

### uFINE array positioning and insertion

The positioning and insertion of flexible arrays in DBS surgeries and brain tumor resection surgeries utilized different micromanipulators, but the operational protocols were similar.

For DBS surgeries, after a burr hole for DBS lead implantation was created and the dura mater was incised, the implantation of the uFINE array occurred prior to the DBS lead placement. The specific procedure was as follows. The NeuroFortis micromanipulator was installed in the Leksell stereotactic frame. The implantation trajectory of the uFINE closely aligned with the planned trajectory of the DBS lead, though slight adjustments were made to avoid blood vessels or sulci of the cerebral surface. Before inserting the Neuropixels probe, a small superficial incision in the pia was done using an arachnoid surgical knife. Subsequently, the uFINE array replaced the dummy module on the micromanipulator.

Sterile ground and reference needle electrodes were placed in the scalp. A sterile data cable connected the headstage to the StairPlex system. During insertion, the uFINE array was guided into place by the shuttle wire, with an insertion depth of 5-6 mm. Once the target depth was reached, the redundant part of the array prefixed at the substrate was released immediately. The shuttle wires were withdrawn and left suspended approximately 3-5 mm above the brain surface, leaving only the flexible array shanks embedded in the brain tissue. The released redundant part ensured that the uFINE array had sufficient distance when moving with the brain surface, preventing it from being pulled out.

For brain tumor resection surgeries, a custom-designed micromanipulator mounted on a commercial multi-articulated arm (Brainlab Co.) replaced the Leksell stereotactic frame and NeuroFortis for uFINE array implantation. All patients undergoing resection surgery were awakened from anesthesia after the skull and dura mater were opened.

### Electrophysiological recordings

In this study, the headstage used (Shanghai Stairmed Technology Co.) had a linear dynamic range of 10 mVpp and a 10-bit sampling resolution. The recording sampling rate was 25 kHz, and impedance measurements were conducted prior to electrophysiological recordings. For some patients participating in the “Word-repeating” task, a set of speakers and a screen were used to provide stimuli, and responses were recorded using a microphone. TTL signals from the task laptop were sent to the digital input channels of the StairPlex system via a multi-function I/O module (NI USB-6009, National Instruments) to synchronize the electrophysiological recordings with behavioral data for each trial.

### “Word-Repeating” task

In this study, three patients performed a “Word-repeating” task. The “Word-repeating” task in this study consisted of two types: Pt.03 was assigned Type I, while Pt.04 and Pt.08 were assigned Type II. The paradigms for the two types of tasks were the same (Figure 4A), but the libraries were different. In each trial, patients needed to listen to a specific word, which was synchronously displayed on the screen, as the “stimulus”. After a fixed delay of 0.6 s, patients were asked to orally repeat the word as the “response” following a 0.1-s white noise prompt. The library for Type I included “Ji Li (吉利),” “Ji Li (极力),” “Ying Wu (英武),” and “Ying Wu (鹦鹉).” The library for Type II included “Xin Nian Kuai Le (新年快乐),” “Zhu Ni Ping An (祝你平安),” “Zao Ri Kang Fu (早日康复),” and “Qin Lao Yong Gan (勤劳勇敢).” A word was randomly selected from the library for each trial.

Pt.03 had an average stimulus period of 0.51 s (std = 0.01 s) and an average response period of 0.65 s (std = 0.09 s). Pt.04 had an average stimulus period of 0.98 s (std = 0.05 s) and an average response period of 1.04 s (std = 0.09 s). Pt.08 had an average stimulus period of 0.98 s (std = 0.03 s). Due to Pt.08’s slower speaking tempo, the average response period was 3.12 s (std = 1.06 s). The three patients completed 101, 79, and 59 valid trials, respectively. It was confirmed before the surgery that all three patients were familiar with the pronunciation and meaning of all the words in the library. TTL triggers were generated during the task using custom code in Python on the task computer. The starting position of each trial was aligned with the starting position of the triggers. The microphone was turned on at the start of the trial and turned off when the trial terminated. Each trial lasted 6 s.

## QUANTIFICATION AND STATISTICAL ANALYSIS

### Spike sorting

Before automatic spike-sorting, the raw signals underwent the following preprocessing steps:

1. High-pass filtering (250 Hz);
2. Removal of high-impedance channels higher than 4 MΩ;
3. Removal of channels with RMS noise higher than 2 σ. Based on empirical information, the SNR for such channels is very low;
4. Removal of global channel synchronization artifacts: When all channels recorded a signal greater than 100 μV simultaneously, the portions of the signal surrounding the maximum absolute value point that exceeded 100 μV were set to zero for 10 ms before and after that point.

This study utilized Kilosort 4.0^36^ (https://github.com/MouseLand/Kilosort) for spike sorting. The specific parameter settings were as follows:

1. Nblocks = 0. The spike drift of the flexible electrode array was minimal, so there was no need for a drift correction algorithm to improve spike sorting accuracy during spike sorting. Statistical analysis of spike position drift (Figure 3A-C) confirmed that drift correction was unnecessary.
2. In spike detection, both “Th_universal” and “Th_learned” thresholds were set to 8.

Subsequently, experienced electrophysiologists (SW) performed manual curation of the automatically isolated clusters using Phy (https://github.com/cortex-lab/phy), with the following specific steps:

1. Clusters of abnormal waveforms were labeled as noise, primarily including sinusoidal oscillations and spikes with durations less than 0.1 ms.
2. When a cluster could be clearly observed to have separation in principal component (PC) space, it was split into two or more clusters. If the spatial spread of waveforms, amplitude, and Inter-spike-interval (ISI) distribution were highly similar, and merging did not result in a higher ISI violation, the clusters were merged in Phy. The ISI violations refer to the rate of refractory period violations^47^.
3. If the ISI violation was less than 0.2 and the FR was above 0.05 Hz, the cluster was labeled as a good unit. Otherwise, the cluster was labeled as multi-units (MUA).

Among the “good units”, those with an average amplitude exceeding 30 μV were considered single units and included in subsequent statistical analyses.

### Assessment of recording quality

First, we calculated the single-unit yield per channel to evaluate the efficiency of electrode recordings in capturing neuronal signals. The yield for each patient was defined as the number of sorted single units divided by the number of usable channels, excluding channels with impedance >4 MΩ or abnormal RMS values (as previously described).

Next, to assess signal quality, we analyzed the amplitude, firing rate, SNR, ACG rise time, trough-to-peak duration, and peak-trough ratio of all sorted neurons. For each metric, the 25th, 50th, and 75th percentiles were calculated to summarize variability and central tendency, as shown in Figure 2E. The specific calculation methods for these metrics are as follows:

1. Peak-trough ratio: Defined as the maximum value among the first and second peak-to-trough ratios in the mean waveform of each unit, as illustrated in Figure 2F.
2. Trough-to-peak duration: Defined as the time interval (ms) between the waveform trough and the subsequent peak (the peak utilized for calculating the peak-trough ratio) in the mean waveform, as illustrated in Figure 2F.
3. SNR: The signal-to-noise ratio (SNR) was calculated as the peak-to-trough amplitude of the mean waveform divided by twice the standard deviation of the data stream.
4. ACG rise time: Computed using custom scripts based on the method described in Wu et.al^48^. The score quantifies temporal characteristics of single-unit activity distributions by fitting a triple-exponential equation to the ACG.

### Waveform classification

Waveforms of single units were extracted and manually curated using custom code, followed by classification using a standardized approach to categorize the units into regular-spiking (RS), triphasic-spiking (TS), and positive-spiking (PS) types, based on the spike waveform recorded at the channel exhibiting the largest amplitude for each unit^49,50^.

As illustrated in Figure 2F, PS (positive-spiking) waveforms were characterized by positive peaks whose magnitude exceeds that of the negative troughs. TS (triphasic-spiking) waveforms display a prominent initial positive peak, followed by a larger negative trough (with a first peak-trough ratio >0.1), and a subsequent positive peak occurring within 1 ms after the initial peak. In contrast, RS (regular-spiking) waveforms typically exhibit dominant negative troughs.

### Assessment of recording stability

#### (1) Spike positions calculation

The spike positions for each single unit were obtained from the “spike_positions.npy” file automatically generated by Kilosort4. Since the uFINE array’s recording sites are arranged in a vertical linear configuration, the file contains only longitudinal position data.

#### (2) Waveform similarity

We assessed recording stability by analyzing the waveform similarity of single units across different recording periods^51^. For each single unit, the mean waveform was calculated within each recording segment. Then, the Pearson correlation coefficients were computed for pairs of average waveform vectors across segments to evaluate waveform similarity.

#### (3) Waveform shape stability in PC space

We assessed waveform shape stability using PCA. We projected all spikes from each single unit into a three-dimensional coordinate system. The waveform length for each spike was set to 2.5 ms. The x-axis and y-axis represented the first and second dimensions of the PCA of all waveforms, while the z-axis indicated the recording length in seconds.

#### (4) The definition of active units

A unit was defined as an “active unit” when both conditions met:

1. Its firing rate was higher than 0.1 Hz;
2. At least one spike fired within each time window.

We standardized the recording time for each patient by dividing their recordings into 10 equal time windows, irrespective of the varying recording lengths.

#### (5) Recording quality of active units in 10 time windows

For each time window, the mean and standard deviation of spike amplitude, trough-to-peak duration, and peak-trough ratio of all active units were calculated. These metrics were used to assess changes in unit waveform characteristics over time, with smaller waveform variations indicating more stable recording quality. The mean firing rate within each time window was used to characterize neuronal activity over time.

To further assess recording stability, we performed ANOVA to compare each parameter across time windows. The p-value for the comparison between each time window with respect to the rest of the windows was calculated, with p > 0.05 indicating no significant difference. The results are shown in Table S2.

#### (6) The estimation of brain pulsation movement

We used custom MATLAB code to extract markers from videos recorded with a surgical microscope including brain pulsation trajectories to estimate the patients’ brain pulsation movement. All microscope videos were recorded at 30 frames per second. A total of six patients had microscope videos that effectively captured brain pulsation. The estimation process was as follows:

1. A segment of video of at least 30 s was extracted, ensuring that both the length calibration marker (typically the cross-section of the silicon substrate holding the uFINE array, with a constant length of 4 mm) and the brain movement marker (usually a clearly visible reflection point corresponding to the pulsation trajectory at the surface of the brain) were unobstructed, and that the shooting angle and distance remained unchanged.
2. The number of pixel points for the length calibration marker was measured, and the length-to-pixel ratio was calculated based on its actual length.
3. Each frame of the video was extracted and converted to grayscale. Regions greater than the grayscale threshold (with a threshold value of 220) were extracted from a fixed area in each frame as the Region of Interest (ROI), with the center of the ROI designated as the brain movement marker.
4. The trajectory of the brain movement marker within the selected video segment in step (1) was extracted and converted into actual distance using the length-to-pixel ratio.
5. The trajectory curve of the brain movement marker was plotted, and the peak-to-peak motion for each pulsation was calculated, yielding the mean peak-to-peak motion as the average brain pulsation movement distance for each patient.

The limitations of the estimation method used in this study were as follows:

1. The videos lacked markers for depth-of-field calibration;
2. Brain movement described a trajectory in three-dimensional space (primarily represented as vertical movement relative to the uFINE array, but also included some lateral movement), while the video only depicted this three-dimensional motion projected onto a two-dimensional plane at the shooting angle.

### Stimulus and response tuning of single units

The FR tuning curve for each single unit was calculated through the following steps:

1. A commonly used speech analysis tool for phonetics, Paart (Phonetic Science, Amsterdam), was utilized to mark the onsets and offsets of the stimulus and response for all trials.
2. The duration from stimulus onset to stimulus offset for each trial was defined as the stimulus duration, and the trial-averaged stimulus duration was calculated. Similarly, the trial-averaged response duration was obtained.
3. One stimulus period was defined as 0.5 s before the stimulus onset to 0.5 s after the stimulus offset. The response period was defined similarly. Each period was divided into time bins with a length of 50 ms. The number of spikes fired during each time bin was counted and converted into spikes per second (Hz) to obtain the FR for that time bin.
4. If a single unit had fewer than 20 trials and a trial-averaged FR below 5 Hz, it was excluded from the tuning analysis. The remaining single units were used in the following analysis. In Figures 4D, 4H and 4I, the raster plot for each single unit displayed only 15 randomly selected trials.
5. The trial-averaged firing rate (FR) and the standard error of the mean (SEM) of FR for each time bin were calculated for each valid tuning-analysis unit.
6. The FR tuning curve was plotted using the trial-averaged FRs from the time bins. To reflect the overall tuning trend, a smoothing sliding window of 50 ms with an overlap of 25 ms was applied.
7. The average FR in the 0.5 to 0.25 s before the stimulus onset was defined as the FR baseline. Each time bin after the stimulus onset and response onset was subjected to a significance test (Wilcoxon signed-rank test, p<0.05) against the FR baseline to determine whether that time bin had significance. Time bins with a significant increase in FR relative to the baseline FR were defined as significant activation periods, while time bins with a significant decrease were defined as significant suppression periods.
8. When a single unit’s significance period lasted continuously over 150 ms^23^, it was defined as a significant activation or suppression unit.
9. The proportions of significant activation units and significant suppression units were calculated for each patient during the stimulus and response periods.
10. The difference between the baseline FR and the FR for each time bin was defined as △FR. A heatmap displayed a matrix of △FR values, with each row corresponding to a valid tuning-analysis unit and each column corresponding to the △FR for each time bin of that valid tuning-analysis unit. This heatmap was used to compare the fluctuations of △FR between units during the stimulus period or response period. The heatmaps for the response and stimulus were sorted in descending order based on the △FR values during the response period.

